# PREMATURE BIRTH AND CESAREAN SECTION ARE ASSOCIATED WITH SIGNIFICANT DIFFERENCES IN NEONATAL CD4^+^ T CELL GENE EXPRESSION AND FUNCTION

**DOI:** 10.64898/2026.02.21.707199

**Authors:** Carlos Jesús Ventura-Martínez, Linda Aimara Kempis-Calanis, Sebastian Mijares-Guevara, Alejandra Cedillo-Baños, Ingrid Yaritzit Carreón-Cortés, Darely Yarazeth Gutiérrez Reyna, Stephania Vázquez Rodríguez, Addy Cecilia Helguera-Repetto, Claudine Irles, Salvatore Spicuglia, Otoniel Rodríguez-Jorge, María Angélica Santana

## Abstract

Premature birth and cesarean section are associated with increased morbidity and inflammatory diseases. However, their impact on neonatal immunity remains incompletely defined. To explore how gestational age and mode of delivery contribute to early immune programming, we analyzed CD4^+^ T cells, central regulators of adaptive responses, from preterm neonates and full-term neonates born by cesarean section or vaginal delivery.

We performed transcriptomic profiling (mRNA-seq) and functional assessment of T cell activation, proliferation, and cytokine production following stimulation. The mode of delivery emerged as a key factor for CD4^+^ T cell transcriptome and function. CD4^+^ T cells from full-term neonates born by vaginal delivery exhibited an immune activation signature, produced higher levels of multiple cytokines, and showed reduced proliferative capacity.

In contrast, prematurity was associated with modest changes in basal gene expression relative to full-term cesarean section neonates. CD4^+^ T cells from preterm neonates displayed enhanced proliferation and increased secretion of inflammatory cytokines (IL-13, TNFα, IL-6, and IL-17F) upon stimulation, consistent with heightened responsiveness. Collectively, our findings show that CD4^+^ T cells from preterm neonates exhibit augmented inflammatory potential, which becomes more regulated at term. Mode of delivery further contributes to this developmental trajectory: cesarean section is associated with a restrained functional profile, whereas vaginal delivery is associated with a mild immune activation signature and increased responsiveness. These results support a model in which neonatal CD4^+^ T cell trajectories are established during fetal life and further modulated at birth, highlighting the layered influence of perinatal factors on immune development.

**Summary sentence:** Neonatal CD4^+^ T cell trajectories are established during fetal life and further shaped at birth by the mode of delivery, influencing early immune responsiveness.

## 1. Introduction

The neonatal period, comprising the first 28 days of life, is the most vulnerable period for child survival. The neonatal immune system exhibits distinct characteristics and is believed to be programmed during the intrauterine period. This results in neonates having limited innate inflammatory and Th1 immune responses, but predominant Th2, Th17, and tolerant immune responses^1–5^. Neonatal T cells exhibit a limited activation response, biased toward innate immunity^6–9^. We previously published transcriptomic and epigenetic analyses of CD4^+7^ and CD8^+6^ T cells from human neonates, compared with adult naïve T cells, showing distinct functional programming characterized by high proliferation, a highly glycolytic metabolism, an innate-biased response, and low effector functions^10^. Furthermore, early events that occur during gestation, childbirth, and the perinatal period have a significant impact on an individual’s development and health throughout life. The immune system’s development and response are also affected by early life events. Premature birth is one of the major causes of death in early life, causing around 28% of deaths in the population of under-5-year-old children in Mexico^11,12^. In addition to the high rate of fatalities, premature babies often suffer morbidities that leave sequels that impact their health for life, particularly those who are born with a very low weight or before 33 weeks of gestational age (wga)^13,14^. Premature babies are highly susceptible to infections and have limited protective immune responses^2,15,16^. However, their immune system is hyperreactive, often leading to highly inflammatory conditions such as sepsis and necrotizing enterocolitis^17–20^. The causes of prematurity have been previously studied; maternal conditions such as obesity, preeclampsia and intrauterine infections have been associated, while maternal and fetal inflammatory responses contribute to preterm labor^21–25^.

The rate of cesarean section (c-section, cesarean) for babies born under 33 wga is between 45% and 72% worldwide^14^. In addition, many full-term babies are also delivered by cesarean section, often planned surgeries without a medical reason. Worldwide, the percentage of cesarean sections as compared to vaginal birth is 32.5% and has been on the rise^26^. However, in Mexico, 55% of total births and most premature babies are delivered by cesarean section, exceeding the world average^11,27^.

The effects of prematurity and birth type on neonatal development and immune response have been studied, but clear results remain lacking. Some epidemiological studies have shown, however, a predisposition of babies born by cesarean section to a series of metabolic and proinflammatory conditions, such as obesity, diabetes, asthma, and autoimmune diseases^27–31^.

To better understand the impact of birth type and gestational age on the immune system, we evaluated CD4^+^ T cells, as these cells are central to the strength and type of adaptive immune response. Here, we present the evaluation of the transcriptome, cell activation, proliferation, and cytokine secretion of CD4^+^ T cells from preterm (28 to 36 weeks of gestation) newborns born by cesarean section, as well as full-term newborns born by cesarean section (cesarean) or vaginal delivery (natural birth). Our results show a greater difference in the transcriptome between full-term neonates born by cesarean section and those born by vaginal delivery, whereas prematurity exhibits only minor changes relative to their controls. The gene expression profile of natural birth CD4^+^ T cells showed a mild “immune activation” signature. Upon stimulation, CD4^+^ T cells from vaginal delivery produced modest but higher amounts of all cytokines, but proliferated less than those from cesarean section, which indicates a predisposition to respond to stimulation. Of note, CD4^+^ T cells from premature babies showed a remarkably high production of cytokines, especially IL-13, TNF-α, IL-6, and IL-17F, and a high proliferation rate as compared to cells from full-term neonates from cesarean section.

Together, these findings suggest that birth mode is a contributing factor in the functional programming of neonatal CD4^+^ T cells, whereas prematurity plays a role in potent inflammatory responses. This study provides molecular and functional evidence that immune trajectories are programmed during fetal life and further modulated at birth, underscoring the immunological relevance of delivery practices and gestational age in neonatal health.

## 2. Materials and Methods

### Type of study and statistical analysis

This study was conducted in an observational cohort of neonates recruited at birth, as described in Supplementary Table S1. Umbilical cord blood was collected at delivery from three independent groups of neonates: (i) full-term neonates delivered vaginally (VD), (ii) full-term neonates delivered by cesarean section (CS), and (iii) preterm neonates delivered by cesarean section (PT). These groups were defined according to naturally occurring clinical conditions, without investigator intervention. Exclusion criteria were defined *a priori* and included neonatal infection (confirmed by culture or compatible clinical findings), congenital malformations, neonatal fever, low Apgar score, maternal diseases known to affect neonatal immune function, and lack of maternal informed consent.

Cord blood mononuclear cells (CBMCs) and purified CD4^+^ T cells were isolated from each neonate and cultured under basal conditions or stimulated *in vitro* with anti-CD3/CD28 antibodies. Each donor therefore contributed paired measurements (basal and stimulated conditions). Accordingly, the study followed a mixed design comprising one between-subject factor (birth group) and one within-subject factor (stimulation).

Because the birth groups were observational rather than experimentally assigned, the statistical analysis focused on prespecified biologically relevant comparisons rather than all possible pairwise group comparisons. Planned comparisons were defined *a priori* according to the study hypotheses. Specifically, we evaluated (i) the effect of stimulation within each birth group using paired analyses and (ii) differences among birth groups under the same experimental condition. Because the study was observational and the hypotheses were defined *a priori*, the statistical analyses were designed to evaluate prespecified biologically relevant comparisons rather than factorial interaction effects. Given the relatively small sample sizes and the inherent biological variability of primary human neonatal samples, nonparametric statistical methods were used throughout. Comparisons among independent birth groups were performed using the Kruskal–Wallis rank-sum test, whereas paired comparisons between basal and stimulated conditions were evaluated using the Wilcoxon signed-rank test. To account for multiple testing across cytokines, *P-*values were adjusted using the Benjamini–Hochberg false discovery rate (FDR) procedure. Unless otherwise indicated, adjusted *P-*values < 0.05 were considered statistically significant.

### Cell purification and stimulation

Neonatal blood was collected as previously described^7^. Briefly, the blood was collected just after the babies’ birth by puncture of the umbilical cord vein, from births through vaginal delivery or cesarean section at Hospital General José G. Parres (Cuernavaca, Morelos, Mexico), Hospital General de Temixco (Temixco, Morelos, Mexico), Instituto Nacional de Perinatología (Mexico City, Mexico), with full informed consent from the mothers and the ethical approval of the hospital (CONBIOÉTICA-17-CEI-001-20160329). An official agreement (File 1372/1706) was granted by Servicios de Salud Morelos to collect the blood samples in Cuernavaca and by the Instituto Nacional de Perinatología (2018-1-149). Samples were processed immediately to obtain mononuclear cells by density gradient centrifugation using Lymphoprep (Axis-Shield). Then, they were cultured overnight in tissue culture dishes (cat. 353003, Corning) to eliminate adherent cells and obtain cord blood mononuclear cells (CBMCs). To obtain CD4^+^ T cells, CBMCs were mixed with the RosetteSep CD4^+^ T-cell enrichment cocktail (STEMCELL) and 1 mL of cultured erythrocytes from the same blood sample, and then separated by a second density gradient. Cells were cultured at 37 °C and 5% CO_2_ in supplemented RPMI medium (10% Fetal bovine serum, 1% glutamine, 100 U/mL penicillin, and 100 μg/mL streptomycin) (Thermo Fisher Scientific). All samples were at least 93% CD3+/CD4^+^, as determined by flow cytometry, and very similar purities were obtained in all populations (Supplementary figure 1). The purity of our populations of CD4^+^ T cells was evaluated with antibodies (anti-CD4 (20-0048-T100) and anti-CD3 (20-0037-T100); TONBO biosciences). The absence of maternal cell contamination and the unstimulated status were verified by staining for CD45RO and CD69, markers of memory and activated T cells. Cells were stimulated by crosslinking both CD3 and CD28 antibodies (1 μg/mL) with a secondary antibody (Goat Anti-Mouse IgG Antibody (H + L) bs-0296G, Bioss).

All samples from term pregnancies were delivered at the same public hospitals from Secretaría de Salud Morelos (SSM), where the decision to have cesarean delivery was made during labor, and all mothers received the same care. For term deliveries, clinical evaluation was performed at birth, ensuring that samples were obtained exclusively from clinically healthy mothers (at the time of delivery) and neonates. We specifically excluded any cases involving maternal diseases like active infections, or any other medical condition identified upon hospital admission. We also excluded samples from neonates that were sick at birth (fever, sepsis, malformations, etc). On the other hand, samples from premature babies were obtained from the Instituto Nacional de Perinatología, which specializes in multiple birth pregnancies or other high-risk conditions. Some of the mothers who participated in this study received antibiotic treatment before parturition. In this group, we included only cases classified as clinically healthy at the time of delivery, excluding cases with active infections. Preterm cases included in this study were associated with fetal growth restriction rather than maternal illness, however, some of the mothers had previous conditions (preeclampsia, obesity, and gestational diabetes). The characteristics of our neonatal cohort is described in Supplementary Table 1.

### RNA preparation and sequencing

We prepared total RNA using TRIzol reagent (Thermo Fisher Scientific, 15596-018, Invitrogen), as indicated by the manufacturer. RNA integrity was analyzed using a Bioanalyzer nano-RNA-Kit (Thermo Fisher Scientific), and only samples with an RNA Integrity Number (RIN) > 8.0 were included. For each sample, 500 ng of total RNA was used to prepare an RNA-seq library using the Illumina Stranded mRNA Prep Ligation kit, according to the manufacturer’s instructions. The samples were sequenced on the HiSeq-2500 Sequencer (Illumina). Sequencing was performed in paired-end mode, with a read length of 75, and a sequencing depth of 47.04 ± 11.3 × 10^6^ reads per sample.

### Transcriptome analysis

The workflow for transcriptome analysis was previously described by Kempis-Calanis *et al*^*7*^. Briefly, it consisted in: i) the evaluation of the Data quality using FASTQC program^32^; ii) elimination of adaptors and low-quality reads with Trimmomatic program (version 0.39)^33^; iii) reads alignment against the reference genome (GRCh38/hg38), using the STAR 2.4.2^a^ software^34^; iv) visualization of the alignment with IGV^35^; v) obtention of the read counts per gene with HTSeq-count^36^; vi) Identification of the differentially expressed genes using DESeq2^37^ package in R (Bioconductor). All samples were normalized and a design formula that accounted for the experimental groups (design = ~ group) was used. Differential expression analyses were then performed using the contrasts of interest: (1) term cesarean section (CS) versus term vaginal delivery (VD), (2) preterm cesarean section (PT) versus term cesarean section (CS). Differentially expressed genes (DEGs) were identified using our predefined statistical criteria (|log2FoldChange| ≥ 1 and FDR < 0.05); vii) identification of the enriched pathways with ClusterProfiler^38^ package in R, using the function EnrichKEGG; viii) visualization of the enriched pathways (enrichplot (Bioconductor)). We also performed an analysis of transcription factor activity based on the expression of their targets using decoupleR^39^ package in R. Additionally, gene set enrichment analysis (GSEA)^40^ was performed in R using the whole DESeq2 results table and the clusterProfiler package. Finally, a protein-protein interaction (PPI) network was generated for the differentially expressed genes (DEGs) with the STRINGdb^41^ R package (v2.22.0), recovering all available interactions from the STRING database (v12.0) for Homo sapiens (taxonomy ID: 9606). Interactions were filtered using a minimum score of 0.7 (high confidence), and isolated nodes were removed from the final network. The network was visualized and analyzed in Cytoscape (v3.10.4)^42^. Functional enrichment analysis for Gene Ontology (GO) Biological Process terms was performed using the STRING enrichment function implemented in the stringApp plug-in for Cytoscape (v2.2.0)^43^ to finalize network annotation.

Note: The anonymized RNA-seq datasets are deposited in the GEO repository (GSE322854).

### Cell activation analysis

Cell response was measured by evaluation of i) the activation markers CD69 and CD25, and iii) AP-1 and NF-κB activation. For the evaluation of activation markers, we used anti-CD69-FITC (cat. 310904, BioLegend) and anti-CD25-APC (cat. 20-0259-T100, Cytek). After 72 hours of stimulation, purified CD4^+^ T cells were collected and incubated with a cell viability reagent (Invitrogen) to assess viability, as well as with anti-CD69-FITC and anti-CD25-APC for 30 minutes at 4°C. After incubation, cells were washed with PBS containing 2% FBS, then fixed with 1% formaldehyde.

To assess transcription factor activation, we evaluated p65 and c-Jun phosphorylation by intracellular staining. Activation of p65 and c-Jun was assessed by evaluation of phosphorylation at specific residues. We used anti–phospho c-Jun (Ser63, cat. GTX61170, GeneTex), secondary anti–rabbit FITC-conjugated antibody (GeneTex), and anti–phospho p65 (Ser529)–APC (Miltenyi Biotec). After cell stimulation, cord blood mononuclear cells (CBMCs) were collected and washed with PBS containing 2% FBS, then washed again with 1X PBS. Subsequently, CD4^+^ T cells were stained with anti-CD4-BV605 (cat. 317438, Biolegend) and a viability reagent (cat. L34975, Invitrogen), and incubated for 30 minutes at 4°C. After incubation, cells were washed with PBS supplemented with 2% FBS and then subjected to a permeabilization protocol, which included fixation with 1.5% formaldehyde for 10 minutes, permeabilization with cold absolute methanol for 10 minutes (0°C or −80°C), a wash with PBS supplemented with 2% FBS, incubation with the antibody solution for 30 minutes at 4°C, and a final wash followed by fixation with 1% formaldehyde. Samples were acquired on an Attune NxT cytometer (Invitrogen), and data were analyzed using FlowJo software (Tree Star, CA). Differences in the levels of activated proteins between conditions were assessed using Mann–Whitney or Wilcoxon tests in GraphPad Prism 7.

### Proliferation assay

CBMCs were left untreated or activated through the TCR and CD28 molecules for 96 h in the presence of CFSE (cat. 21888, Sigma) 50 nM. CD4^+^ T cells were followed by staining the CD4 molecule (CD4-PerCP-Cyanine 5.5 (cat. 65-0048-T100, TONBO)). Cell division was evaluated by CFSE dilution using an Attune cytometer (Life Technologies) and analyzed with FlowJo X software. The percentage of divided cells was used for the statistical analysis. This represents the frequency of cells within the precursor population, or the fraction of the original population that underwent at least one division.

### Soluble cytokines/molecules

Production of cytokines and other soluble molecules by CD4^+^ T cells (2 × 10^6^ cells/mL) stimulated for 72h through the TCR and CD28 was evaluated using the Legendplex kit (cat. 741028, Biolegend), according to the manufacturer’s instructions. The following cytokines were evaluated: Th1 (IFN-γ and TNF-α), Th2 (IL-13, IL-5, IL-6, and IL-4), Th9 (IL-9), Th17 (IL-17A and IL1-7F), Th22 (IL-22), Treg (IL-10), and IL-2 with the standard Th kit. In addition, we evaluated innate cytokines (IL-7, IL-1β, and IL-8), TGF-β, and cytotoxic molecules (GZMB and Perforin) relevant to neonatal cells using an ad hoc kit. Cytokine quantification was performed by flow cytometry using an Attune cytometer (Life Technologies).

## 3. Results

### The type of birth is associated with a significant change in the gene expression profile

To investigate the impact of preterm birth and cesarean section on the gene expression profile of CD4^+^ T cells, we obtained transcriptomes of CD4^+^ T cells from preterm neonates born by cesarean section and full-term newborns born by cesarean section or vaginal delivery (VD). First, we purified cells from preterm newborns (28 to 33 wga) born by cesarean section (preterm, PT) and full-term babies (37 to 41 wga) born by cesarean section as controls (cesarean, CS). Additionally, we purified cells from neonates born full-term via vaginal delivery (VD) to compare them with those from term cesarean section. To ensure cell purity and prevent cross-contamination with maternal blood, we evaluated CD45RO and CD69 expression in each sample by flow cytometry. As shown in Supplementary Figure 1, all cells showed comparable purity across populations. All samples had CD3+CD4^+^ purity above 93% and were negative for CD45RO^hi^ or CD69 expression. The transcriptome (Supplementary Tables 2-5) was analyzed as previously described^7^. As an indication of the dispersion of the data, we show a principal components graph (Fig. 1A). To our surprise, the biggest separation among the three different CD4^+^ T cell populations was found between CD4^+^ T cells obtained from neonates born at term by cesarean section (CS) and those of vaginal delivery (VD). Preterm (PT) samples were dispersed, perhaps because the ages varied between 28 and 33 wga. Preterm and full-term babies born by cesarean section showed no real separation in the PCA. We present a volcano plot of differentially expressed genes (DEGs) between vaginal delivery and cesarean section samples, showing 600 differentially expressed genes (Fig. 1B). Some of the genes overexpressed by the natural birth cells were *IL1B, TNFA, CD40, LTBR, CD36, FOSB*, and *FGR*, which are involved in immune responses (Figure 1B). On the other hand, cells from cesarean sections expressed KLF9, MIR186, and MIR1244-4, all of which are associated with proliferation/differentiation. We used the overexpressed genes from these two populations to perform overrepresentation analysis with clusterProfiler^38^. Only one pathway, taste transduction, was found enriched in the full-term cesarean section samples as compared to natural birth samples. This path contains genes from G protein signaling pathways with innate immune functions^44,45^. In the VD samples, however, 27 pathways were found enriched (Supplementary Fig. 2A). The most significant pathways and their corresponding genes are shown in Figure 1C-D. These pathways are related to cytokine and chemokine expression, as well as their receptors, particularly those associated with inflammatory pathways such as TNF and IL-1; T cell activation; signal transduction (CD40, ICAM1, MPK, and FOSB), and cell motility (amoebiasis). Next, we identified differentially expressed genes between CD4^+^ T cells from preterm (PT) and full-term (CS) neonates, both from cesarean sections (Supplementary Fig. 2B). Only 16 genes were found differentially expressed, with the overexpression of CLIC3, a chlorine channel involved in pH, volume, and cell growth regulation, PTPRN2, a phosphatase involved in the secretory pathway, and PTGDR2 involved in prostaglandin metabolism. Overrepresentation analysis didn’t identify any pathways enriched with these DEGs. All the DEGs from the 2 comparisons are shown in Supplementary Fig. 3.

**Figure 1.**
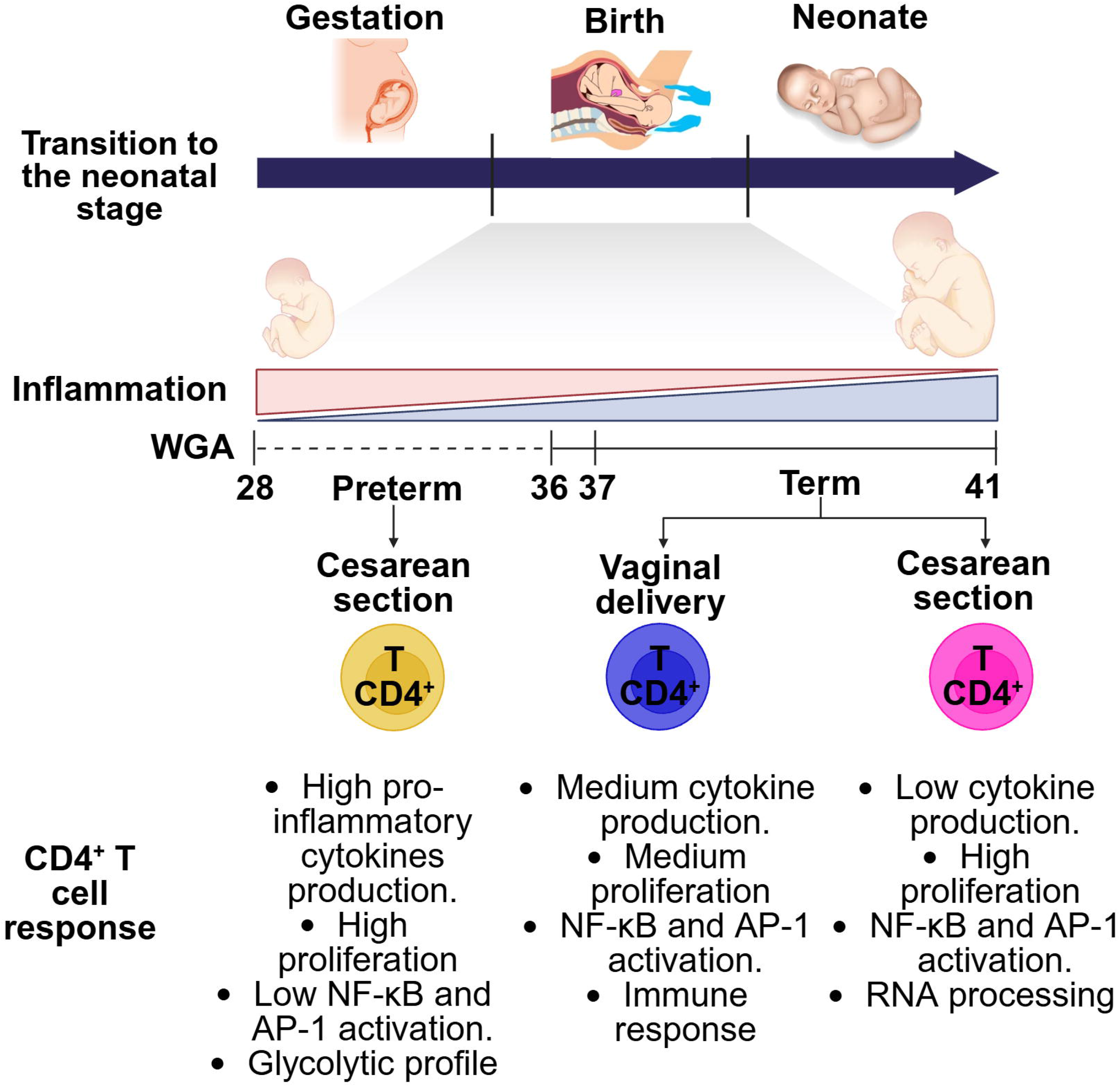
Transcriptomic profiling of CD4^+^ T cells from neonates born at term by vaginal delivery and cesarean section, or preterm neonates. **(A)** Principal component analysis of CD4^+^ T cells from neonates born by vaginal delivery (VD) or cesarean section (CS) at term, and preterm (PT). **(B)** Volcano plot showing the differentially expressed genes between vaginal delivery and cesarean section samples. **(C)** Pathway enrichment by over-representation analysis with clusterProfiler between vaginal delivery and cesarean section samples. **(D)** Network of genes and pathways of the vaginal delivery condition.

We also performed gene set enrichment analysis to further identify processes and pathways potentially relevant to each of the three conditions. In the comparison between VD and CS samples, we identified Biological Processes and Pathways related to adaptive immune response, effector response, chemotaxis, and cytokine production in vaginal delivery CD4^+^ T cells, while CS transcriptomes were enriched for pathways of RNA processing and splicing (Fig. 2A-B and Supplementary Fig. 4). The protein-protein interaction network generated with these genes is shown in Fig. 2C, which shows two main subnetworks: the bigger one related to immune response (*TNF, CD36, CD40, ICAM1, NKG7, GNLY, IL-1β, NLRP3, EOMES, TBX21, PTGS2, BCL6*, and others) and the smaller one related to regulation of cell cycle (*FOXM1, CDC45, MCM10, CDC20, TICRR, AURKB*, and others). When comparing PT and CS samples, a low oxygen response was observed in the PT samples (Fig. 2), as well as a notable enrichment of pathways related to proliferation, glycolysis, aminoacid biosynthesis, and the cytoskeleton (Supplementary Fig. 4).

**Figure 2.**
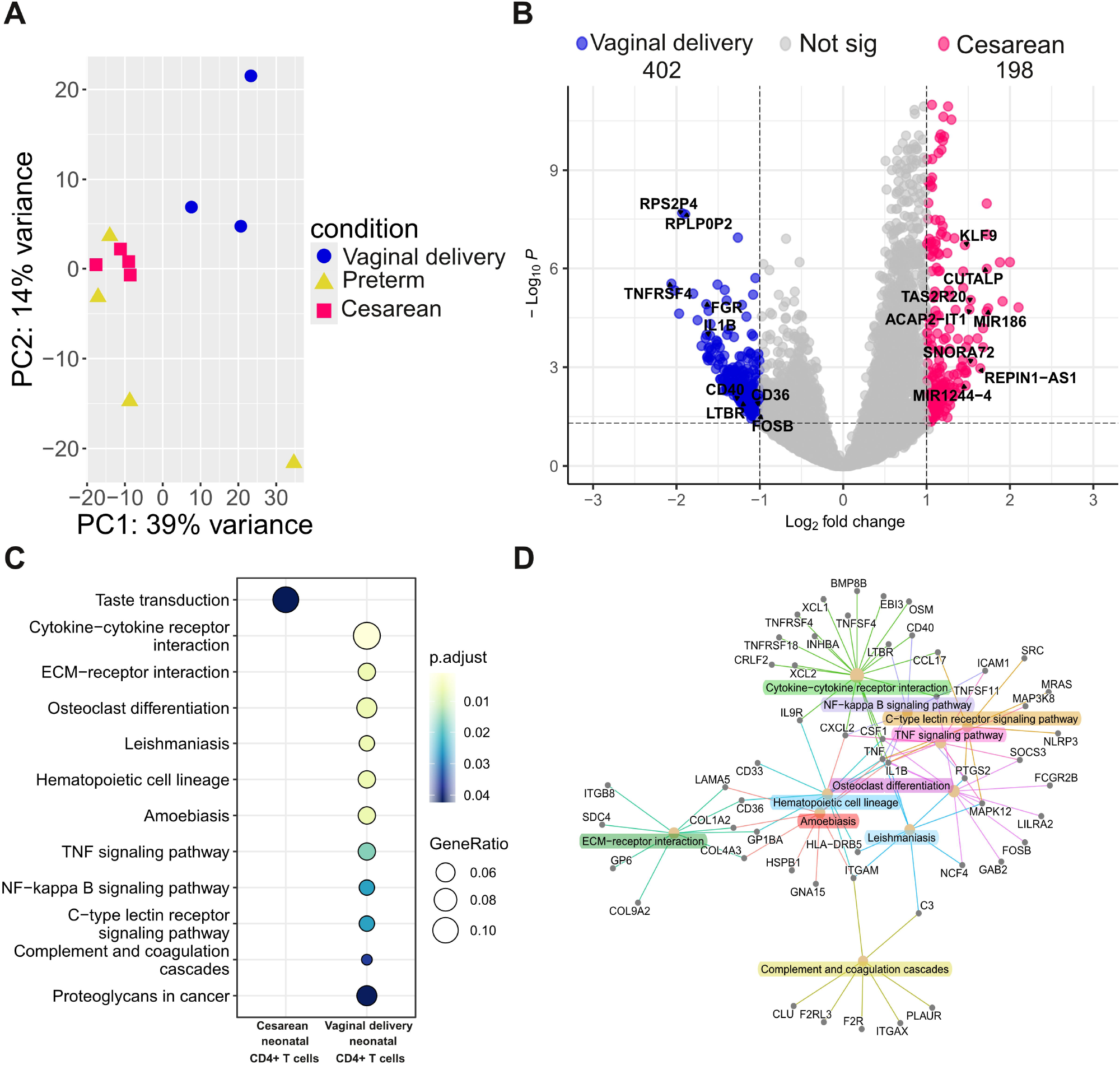
Gene set enrichment analysis and protein–protein interaction (PPI) network. **(A)** Gene set enrichment analysis of CD4^+^ T cells from full-term neonates born by vaginal delivery (VD) or cesarean section (CS), or preterm cells **(B)**, using the GO Biological Processes databases. **(C)** Protein–protein interaction (PPI) network of differentially expressed genes in CD4^+^ T cells from VD and CS neonates constructed using the STRING database and annotated with the most representative GO Biological Processes terms.

To identify transcription factors potentially responsible for the observed transcriptomic signatures, we performed Transcription Factor Activity analysis with decoupleR^39^. In Figure 3, we show the identified transcription factors (TF). When comparing VD and CS samples, we identified NF-κB, AP-1, STAT1, 3, 6, 5A, HIF1A, and MYC activity in CD4^+^ T cells from vaginal delivery, which are typically associated with inflammatory responses.

**Figure 3.**
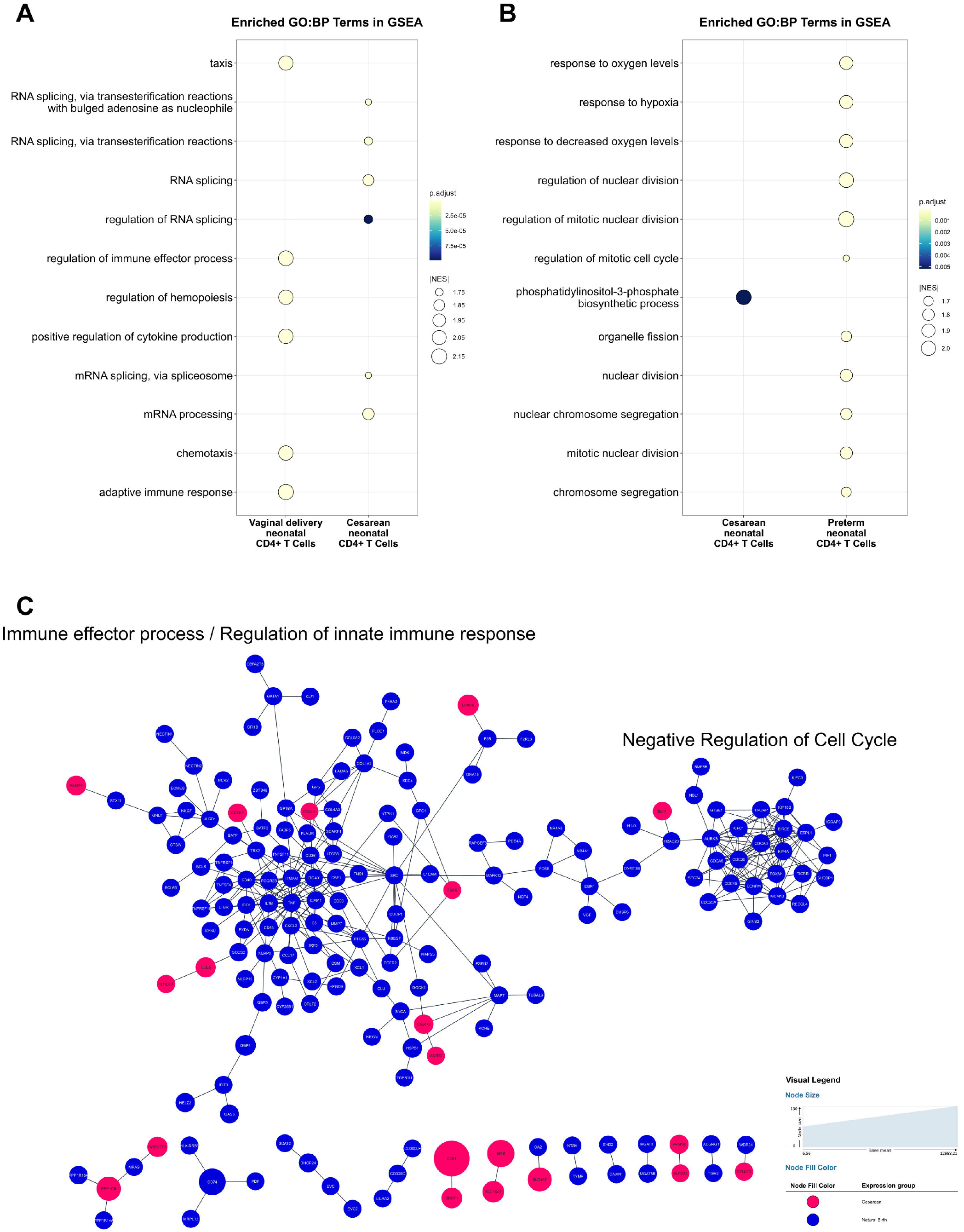
Transcription factor activity in neonatal CD4^+^ T cells. **(A)** Heatmap showing inferred transcription factor activity scores (z-scores) for selected transcription factors across individual samples from neonates born by vaginal delivery (VD), cesarean section (CS), and preterm birth (PT). Colors indicate relative activity levels, ranging from low (blue) to high (red). Samples and transcription factors were organized using unsupervised hierarchical clustering. **(B)** Comparative analysis of transcription factor activity between vaginal delivery and cesarean section. Bars represent differential activity scores, ordered from lowest to highest, with color indicating the associated p-value. **(C)** Comparative analysis of transcription factor activity between preterm birth and cesarean section, displayed analogously to panel B.

These results expand our previous report, which showed that, compared with adult CD4^+^ T cells, neonatal cells from vaginal delivery exhibit a gene expression profile characterized by a limited immune response^7^. However, compared with CS cells, CD4^+^ T cells from VD exhibit an immune activation signature, presumably due to brief activation associated with vaginal delivery, though not reaching adult levels. On the other hand, the transcription factor analysis didn’t identify statistically significant hits in the comparison between CS and PT samples, perhaps due to the small differences in gene expression. Taken together, these results indicate that CD4^+^ T cells from vaginal delivery show a mild “immune activation” genomic signature, while cesarean section cells lack this immune signature.

### Activation of CD4^+^ T cells is different according to the birth type

We next sought to determine whether differences existed in the responsiveness of the three CD4^+^ T cell populations to TCR/CD28 stimulation. Therefore, we stimulated CD4^+^ T cells from full-term neonates born by vaginal delivery or cesarean section, or preterm babies born by cesarean section, in vitro by crosslinking soluble antibodies (CD3/CD28) with an anti-mouse antibody for 72h, and evaluated the expression of the activation markers CD69 and CD25. As shown in Fig. 4A-B, CD4^+^ T cells from all three populations were similarly activated. Interestingly, CD25 expression was higher (23-51%) than CD69 expression (5-13%) in all three populations at that time point (Fig. 4B).

**Figure 4.**
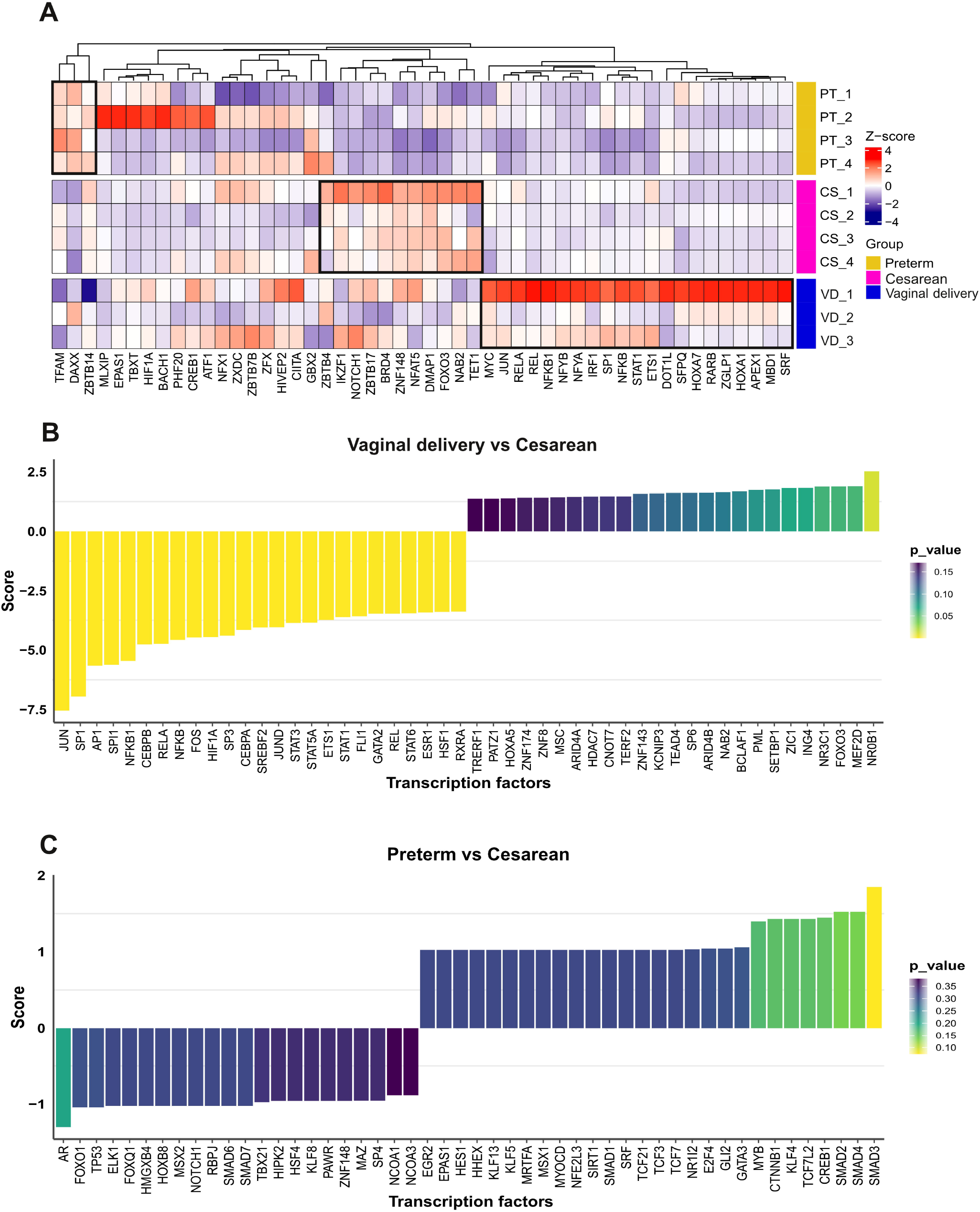
Evaluation of CD69 and CD25 markers and transcription factors activation. **(A-B)** We assessed T cell activation by measuring the expression of CD69 and CD25 on the cell surface. Purified CD4^+^ T cells from term and preterm neonates were either left unstimulated or stimulated under the indicated conditions for 72 hours. The percentages of CD69+ and CD25+ **T** cells were determined by flow cytometry. Mean values are shown above the error bars. **(C-D)** For transcription factor activation, we evaluate phosphorylation of c-Jun and p65 as markers of AP-1 and NF-κB activation, respectively. CBMCs were either left unstimulated or stimulated under the indicated conditions for 90 minutes (to measure c-Jun activation) or 60 minutes (for p65 activation). Mean fluorescence intensity (MFI) values for the indicated phosphoproteins were determined by flow cytometry. **(E-F)** MFI normalized values. The Wilcoxon test was used for paired samples, and the Mann–Whitney U test was used for unpaired samples. \**P* < 0.05, \*\**P* < 0.01, \*\*\**P* < 0.001. For the evaluation of CD69 and CD25 expression, the sample sizes were vaginal delivery = 13, cesarean = 9, and preterm = 3. For the evaluation of transcription factor activation, the sample sizes were: vaginal delivery = 7, cesarean = 8, and preterm = 5.

We also evaluated the phosphorylation of two important transcription factors as a measure of their activation upon TCR/CD28 signals, p-ser529-p65 (classical NF-kB)^46^ and p-ser63-c-Jun (classical AP-1)^47^, as previously reported^48^. The gating strategy for these evaluations is shown in Supplementary Fig. 5A. All three populations showed p65 activation (phosphorylation) upon stimulation (Fig. 4C and E), whereas preterm cells exhibited lower MFI levels (Fig. 4C). However, we were not able to evaluate other members of the NF-kB family. On the other hand, a higher induction of c-Jun activation (phosphorylation) was observed in the samples from vaginal delivery as compared to both term and preterm neonates born by cesarean section (Fig. 4D-F). However, the cesarean section term CD4^+^ T cells showed lower activation of c-Jun upon stimulation, while preterm cells showed overall lower activation (Fig. 4F). This is in agreement with the higher expression of MAPKs and c-Fos in the natural birth samples in the transcriptomic analysis (Fig. 1D), as well as with the high activity of Jun and Fos identified in the Transcription factor activity analysis (Fig. 3). Of note, it has been reported that heightened Calcium signaling, but lower NFAT activation and NFAT-mediated transcription, occur in neonatal T cells^49,50^. In this regard, it has been described that activation of NFAT (calcium signals) without proper activation of AP-1 signals can lead to the activation of the Th2 cytokine profile, characterized by a higher expression of IL-4 and IL-13, and predisposing to allergies and asthma, as has been observed in the cesarean section-born neonates^51^.

### T cells from preterm and full-term neonates born by cesarean section show high proliferation rates

We evaluated the proliferation of the three CD4^+^ T cell populations in response to activation by soluble antibodies (CD3/CD28). The gating strategy used for this is shown in Supplementary Fig. 5B. In response to stimulation by TCR/CD28 signals, all the populations proliferated for up to three or four generations (Fig. 5A). A higher percentage of cell division was observed in CD4^+^ T cells from cesarean section babies, both full-term and preterm (Fig. 5B). This is consistent with the gene expression profile and functional annotation of preterm CD4^+^ T cells (Fig. 2), which indicate cell cycle progression. This high activation-induced proliferation in preterm and cesarean cells contrasts with the results for transcription factor activation, particularly for c-Jun, which was more pronounced in vaginal delivery CD4^+^ T cells but did not reach the same levels in preterm cells.

**Figure 5.**
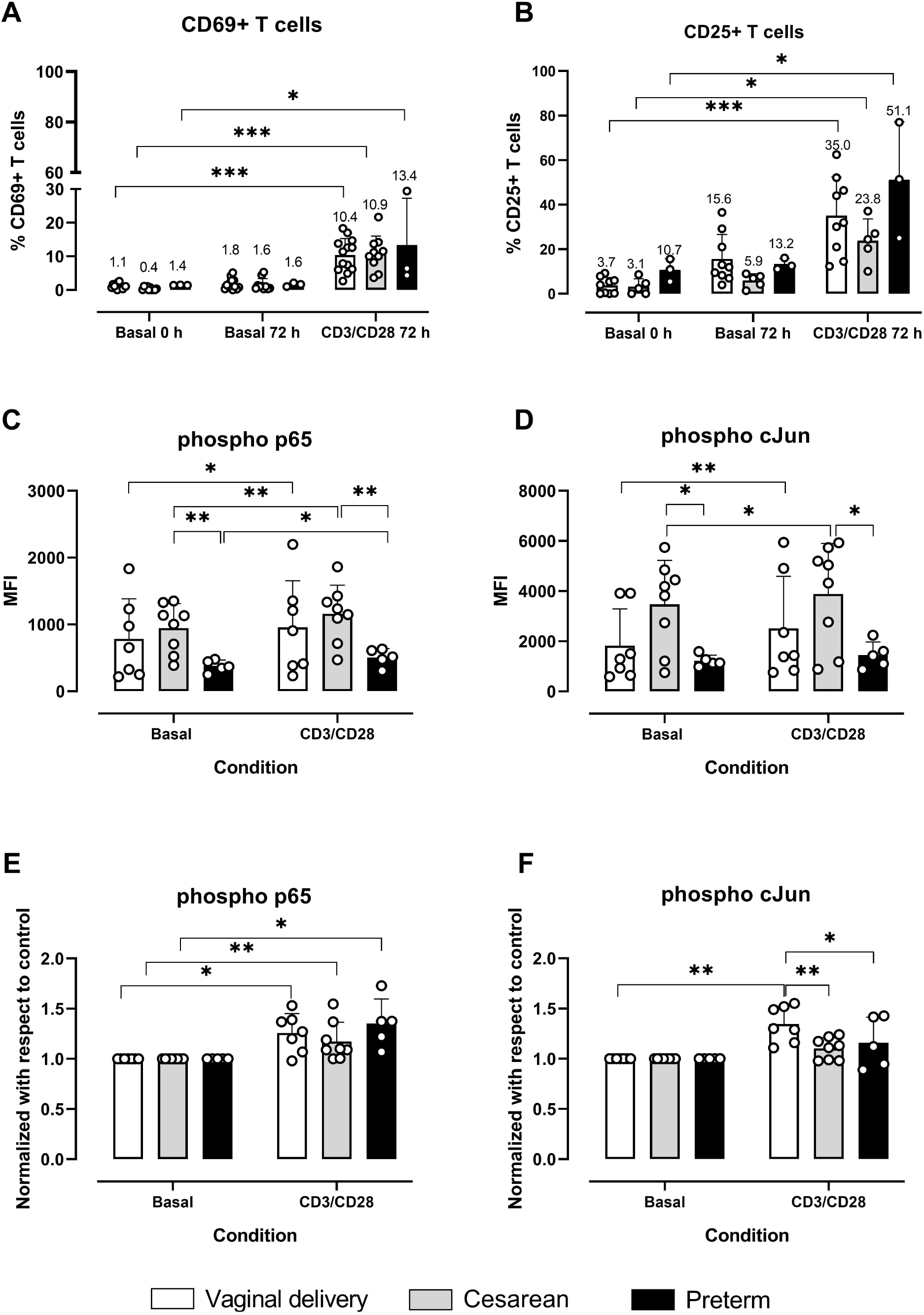
Proliferation assay. CBMCs from term and preterm neonates were stained with CFSE and either left unstimulated or stimulated by crosslinking CD3 and CD28 molecules for 96 hours, followed by staining with anti-CD4–PerCP-Cy5.5. **(A)** Representative histograms of CFSE dilution in vaginal delivery, cesarean section, and preterm birth samples; **(B)** Percentage of division in unstimulated or stimulated CD4^+^ T cells. Statistical comparisons were performed using the Wilcoxon test for paired samples and the Mann– Whitney U test for unpaired samples. \**P* < 0.05. The sample sizes were as follows: vaginal delivery = 5, cesarean = 6, preterm = 5.

### CD4^+^ T cells produce cytokines from all Th phenotypes when stimulated in isolation

To evaluate the functional properties of the three neonatal CD4^+^ T cell subsets, cells were stimulated through TCR/CD28 for 72 h. Cytokines representative of major T helper (Th) lineages were then quantified in culture supernatants using a LegendPlex multiplex assay and analyzed by flow cytometry. Upon TCR/CD28 stimulation, CD4^+^ T cells from all three populations produced IL-2 (Fig. 6). However, average production was highest in preterm cells, followed by those from vaginal delivery, and only a small production was detected in cesarean section cells, as shown in Figs. 6 and 7.

**Figure 6.**
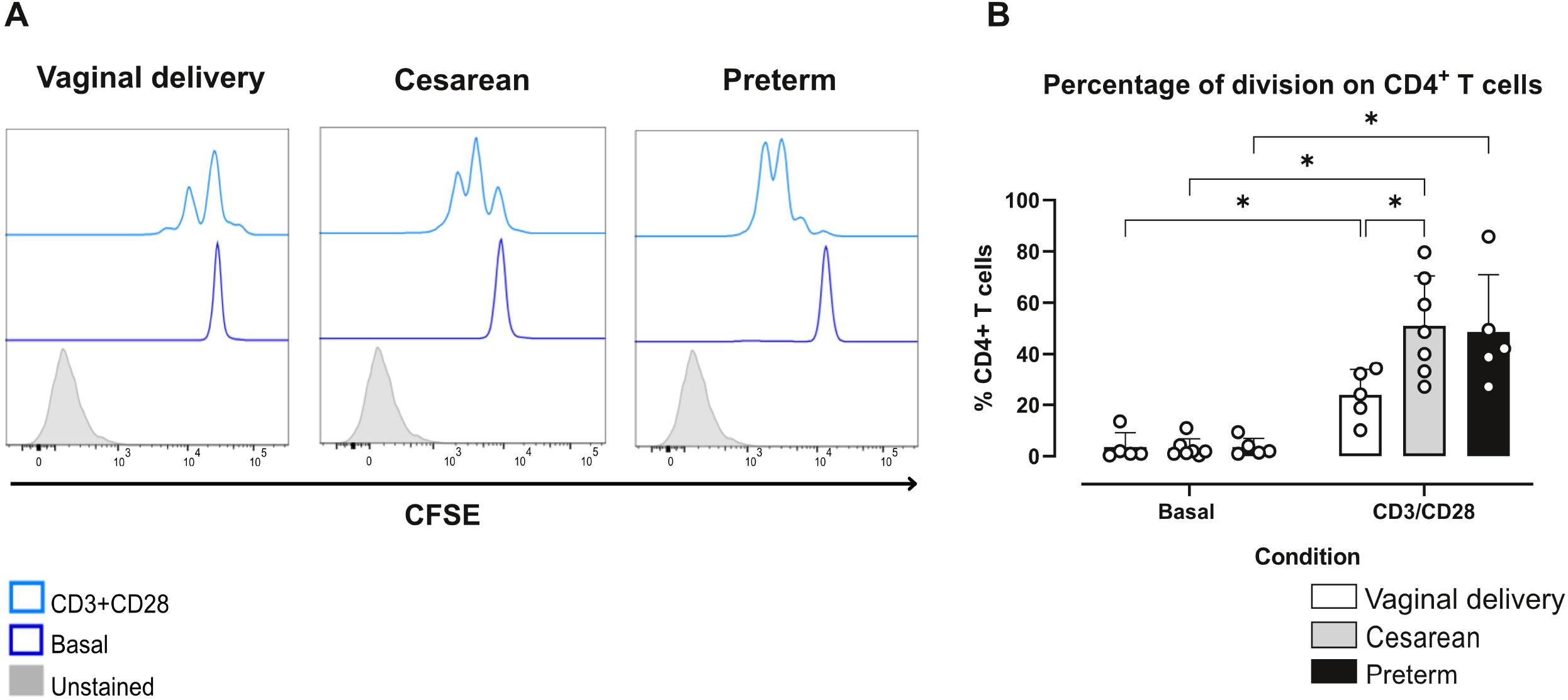
T helper cytokines, innate cytokines, and cytotoxic molecules produced by CD4 ^+^ T cells. Purified CD4^+^ **T** cells from term and preterm neonates were either left unstimulated or stimulated by crosslinking CD3 and CD28 molecules for 72 hours. Cytokines were measured in the supernatants of cultured cells. Cytokine concentrations were analyzed using the Wilcoxon test for paired samples and the Kruskal–Wallis test for unpaired samples, followed by the original Benjamini–Hochberg (BH) false discovery rate (FDR) correction for multiple comparisons. \**P* < 0.05, \*\**P* < 0.01, \*\*\**P* < 0.001. The number of samples were: vaginal delivery = 16, cesarean = 7, and preterm = 4.

**Figure 7.**
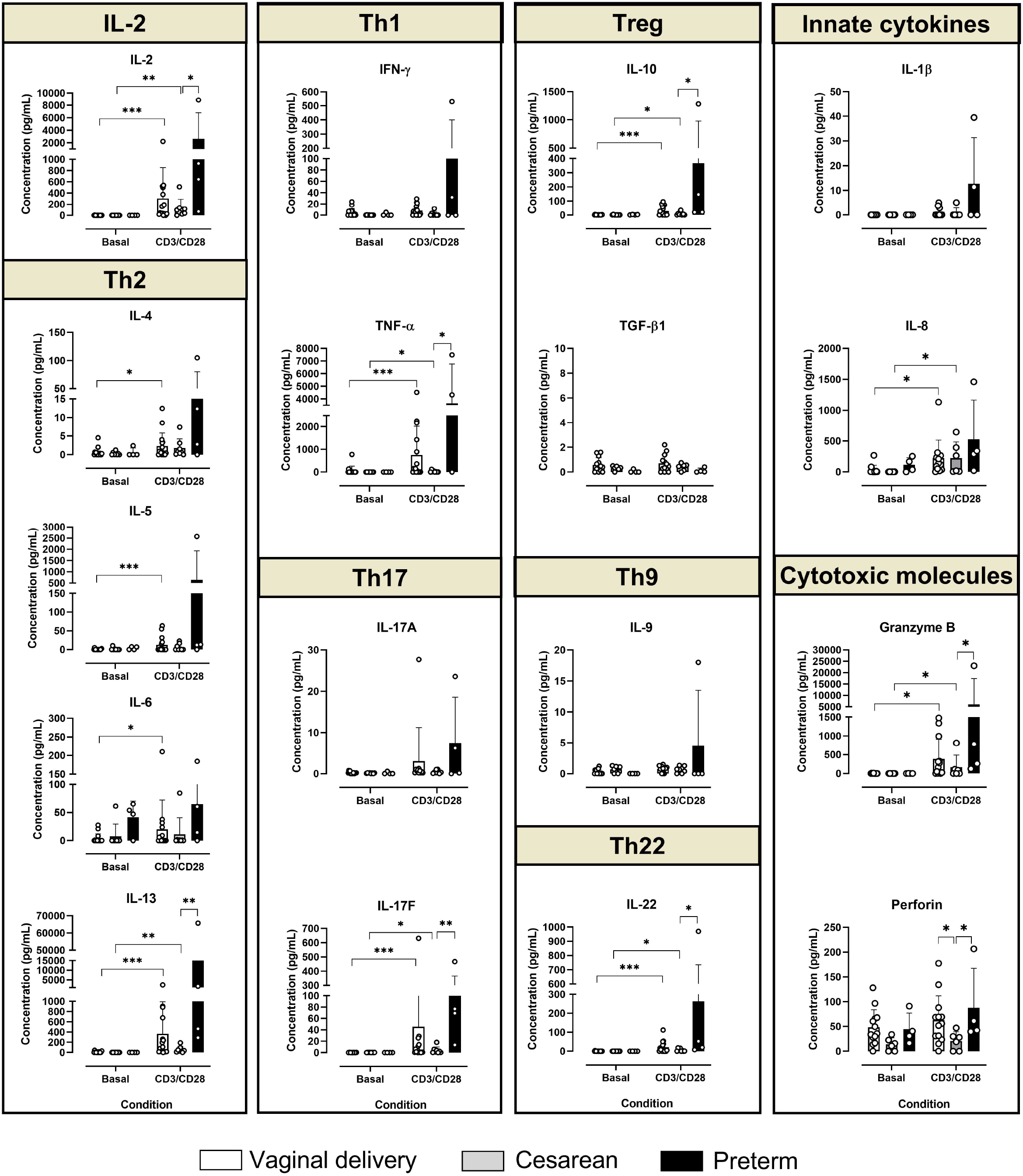
Heatmap of cytokine and cytotoxic molecules production by CD4^+^ T cells. Heatmap showing the normalized concentrations of T helper cytokines (IFN-γ, TNF-α, IL-13, IL-4, IL-5, IL-6, IL-17A, IL-17F, IL-22, IL-9, IL-10, TGF-β1 and IL-2), innate cytokines (IL-8 and IL-1β), and cytotoxic molecules (Granzyme B and Perforin) produced by CD4^+^ T cells from neonates born by vaginal delivery (VD), cesarean section (CS), or preterm (PT) under basal or stimulated (CD3/CD28) conditions. Cytokine concentrations were normalized by row (Z-score of log10(concentration + 1)). Columns represent individual donors, while rows correspond to each cytokine or effector molecule. The top annotations indicate the experimental condition (Basal or CD3/CD28) and the study group (VD, CS, or PT). Color intensity reflects relative concentration levels, with blue indicating higher and yellow lower values. The number of samples were: vaginal delivery = 16, cesarean = 7, and preterm = 4.

CD4^+^ T cells from vaginal delivery produced a significant amount of all four Th2 cytokines (IL-4, IL-5, IL-6, and IL-13). Cells obtained from the full-term cesarean section neonates only induced a small amount of IL-13. On the other hand, cells from preterm neonates showed higher expression of all Th2 cytokines, with markedly elevated IL-13 production. The evaluation of Th1 cytokines showed, as expected, no significant induction of IFN-γ upon stimulation, yet, within the low amount of this cytokine, a higher level was observed in the cells of vaginal delivered neonates compared with those from cesarean section. Again, a high degree of dispersion was observed in preterm neonates, with some samples showing higher IFN-γ levels. TNF-α expression was observed in full-term babies, with lower levels in cesarean section neonates. The cells of the preterm babies showed a very high expression of this cytokine.

From the Th17 profile, only IL-17F showed a significant induction by TCR/CD28 signals in full-term neonates. However, the expression of this cytokine was significantly higher in preterm cells than in term cells from cesarean section neonates.

Regarding the Treg phenotype, IL-10 was expressed in all stimulated cells, with a tendency to higher expression in preterm cells, while TGF-β1 was not detected.

On the other hand, IL-22 was induced by TCR/CD28 signals in all cell populations, but was higher in preterm babies than in cesarean section neonates. Finally, IL-9 expression was very low in all samples.

Other molecules relevant to neonatal cell biology were also evaluated, like innate cytokines (IL-1β and IL-8), IL-7, and cytotoxic molecules (GZMB and PRF), which are shown in Fig. 7. Only two samples of pre-term babies induced the expression of IL-1β. The chemokine IL-8 was produced in all three cell populations of neonatal T cells, as was previously described^52^. Interestingly, Granzyme B expression was stimulated by TCR/CD28 signals, whereas perforin expression remained stable, with higher expression in cells from babies born by vaginal delivery at term and pre-term by cesarean section. The biological significance of this expression in neonatal CD4^+^ T cells is yet to be determined.

Although not all comparison reached statistical significance, due to aspects such as high dispersion of the data, present among all groups but especially important in preterm samples, it is clear that stimulation (CD3+CD28) induces the production of all cytokines, which is stronger in PT samples, then VD samples, and barely detectable in CS samples, as shown in Fig. 7. Also, mean expression of all cytokines and molecules induced is much higher in preterm (PT) samples than Term samples.

To further explore neonatal T cell activation, we evaluated cytokine production upon stimulation in the presence of other mononuclear cells (CBMCs). T cells express a more restricted panel of cytokines and molecules (Supplementary Fig. 6). Of note, only IL-6, IL-10, IL-1β, and IL-8 are produced in all three populations, as well as Granzyme B and perforin. However, IL-1β is more produced in natural delivery CBMCs, which also produce IL-5. On the other hand, in cells from preterm babies, IL-5, IL-13, IL-17F, and IL-22 are also significantly produced. Besides IL-8, which is the most highly expressed across all three populations, IL-6, IL-13, IL-17F, and Granzyme B are the most highly expressed in preterm CBMCs. The mechanism underlying the discrepancies between the cytokines and molecules produced by CD4^+^ T cells in isolation and those produced in the presence of other CBMCs is unclear, but one possible explanation is the direct or indirect roles of the other cells present in the CBMC culture, such as CD8^+^ T cells.

## 4. Discussion

The transcriptomic results identified a genomic signature of CD4^+^ T cells from vaginal delivery (VD) compared with cesarean section (CS) in term neonates. The gene expression profile was enriched for pathways and processes associated with immune responses, including T cell activation, cytokine pathways (TNF, IL1, and IL17), and Integrin and adhesion molecule signaling. We call this feature of CD4^+^ T cells the “vaginal delivery-associated immune activation” signature, which is linked to a greater response to TCR/CD28 stimulation *in vitro*. However, it remains unclear whether this immune activation signature is an effect of parturition, indicates neonatal T cells’ participation in labor induction, or both, or is attributable to other confounding factors that cannot be excluded. One possible hypothesis is bystander activation of CD4^+^ T cells during labor, driven by signals that promote parturition, such as maternal cytokines^53,54^. However, other hypotheses, such as stress molecules from passage through the birth canal or longer-term changes in oxygen levels, among others, cannot be excluded. Mothers of all full-term neonates were from the same hospital, were clinically healthy, and received no pharmacological treatments before birth. On the other hand, most mothers of preterm newborns received antibiotic treatment, and some presented with obesity or preeclampsia, while one mother had gestational diabetes. Of note, at the moment of delivery, there weren’t symptoms or diagnostic of infection or inflammatory conditions for any of the mothers or the newborns participating in the study. Although it is possible that the results from preterm samples were affected by the conditions of the mothers, it is unlikely. The transcriptome of pre-term samples clustered with that of full-term cesarean section babies. Furthermore, pre-term birth samples didn’t express CD45RO or CD69 before stimulation, but sowed strong proliferation and a high production of pro-inflammatory cytokines upon activation.

Following *in vitro* stimulation with soluble antibodies (CD3/CD28), CD4^+^ T cells from all three populations showed similar levels of activation markers (CD25 and CD69) at 72h. Notably, CD69 levels were low (5-13%) in neonatal T cells, while CD25 was more expressed (25-51%) in all three populations upon stimulation. This indicates that typical T cell activation markers may not function similarly in neonatal cells. Additionally, we evaluated p65 (NF-κB) and c-Jun (AP-1) phosphorylation as a proxy of their activation. Higher activation (phosphorylation) of c-Jun and p65 was observed in VD and CS samples as compared to PT ones. This could affect the gene expression profile of PT CD4^+^ T cells, including genes that require AP-1 or NF-κB activation. Of note, it has been previously reported that neonatal VD CD4^+^ T cells have diminished AP-1-dependent transcription when compared to their adult counterparts^50^. Notably, it has been reported that NFACTc2 expression and NFAT activation in neonatal T cells are also diminished relative to those in adult naïve cells^49,55,56^. However, we cannot exclude the possibility of other AP-1 and/or NF-κB members participating.

Regarding cell proliferation in response to stimulation (TCR/CD28 signals), a higher percentage of cell division was observed in CD4^+^ T cells from CS babies, both full-term and preterm (Fig. 5), as compared to VD cells. This contrasts with c-Jun activation, which is higher in VD samples.

Upon stimulation, purified CD4^+^ T cells from VD produced small but significant amounts of cytokines from phenotypes Th1, Th2, Th17, Th22, Treg, IL-8, IL-2, and cytotoxic molecules granzyme B and perforin (Figs. 6 & 7). However, when stimulated in the CBMC mixture, T cells only produced IL-5, IL-6, IL-1β, IL-8, granzyme B, and perforin. This suggests that full-term neonatal CD4^+^ T cells from VD have great immune potential, which may be modulated by their regulatory environment, either through soluble or cellular factors. Another confounding factor could be the reactivity of the CD8+ T cells present in these mixtures, which may also produce cytokines and other molecules. On the other hand, CD4^+^ T cells from term CS neonates produce very low levels of only a few cytokines, including detectable levels of IL-2, IL-13, TNF-α, IL-17F, IL-10, and IL-22, as well as normal levels of IL-8, granzyme B, and perforin. Of note, CS CD4^+^ T cells produce fewer IFN-γ, TNF-α, IL-1β, and perforin than NB cells. In the CBMCs mixture, CS cells produce only IL-10, IL-6, IL-1β, IL-8, granzyme, and perforin. This might indicate that CS cells are blocked from a Th1 response, while VD cells are more prone to respond in this way, which correlates with T-bet (TBX21) expression in VD cells. This is in agreement with the previously reported predisposition of cesarean section newborns to asthma and allergies^30,57^.

Preterm (PT) cesarean CD4^+^ T cells express a variable quantity of cytokines, with a high average. Due to dispersion, not all data were significant; however, cytokines IL-2, IL-6, IL-17F, TNF-α, and IL-8, as well as the molecules granzyme B and perforin, were produced at high levels by these cells in isolation. This is consistent with previous reports associating IL-6^58^, TNF-α59 and IL-8^60^ in preterm CD4^+^ T cells. Of note, IL-6, IL-13, IL-17F, IL-22, granzyme B, and perforin were produced significantly higher than in term CS CD4^+^ T cells. In the culture of CBMCs, PT T cells produce IL-5, IL-6, IL-13, IL-17F, IL-10, IL-22, IL-1β, IL-8, granzyme B, and perforin. Of those, IL-5 and IL-22 were produced in higher amounts than in CS term T cells. This result confirms that preterm CD4^+^ T cells are highly inflammatory, as previously suggested^58,61^; however, more cytokines and molecules might be relevant in the pathophysiology of preterm labor than previously recognized, and special attention should be paid to the relevance of IL-6, IL-13, TNF-α, and IL-17F in the immune pathologies of the preterm neonates.

Together, our findings suggest that neonatal CD4^+^ T cells acquire the capacity to mount inflammatory responses in utero, before term, as evidenced by the heightened functional profile observed in preterm neonates. By term, these cells appear to transition toward a more regulated or tolerogenic state, with reduced responsiveness, as evidenced by samples obtained from cesarean section deliveries. In contrast, the physiological processes associated with vaginal birth appear to actively influence CD4^+^ T cell reprogramming, promoting a more complex and functionally balanced immune profile, which is responsible for the “immune activation” profile. This vaginal delivery-associated program likely reflects exposure to perinatal environmental cues that fine-tune CD4^+^ T cell responsiveness within the broader cellular and tissue context.

## Acknowledgments and sources of funding

We thank the Transcriptomics and Genomics Marseille-Luminy (TGML) sequencing platform. We thank Hospital General José G. Parres (Cuernavaca, Morelos, Mexico), Hospital General de Temixco (Temixco, Morelos, Mexico), and the Instituto Nacional de Perinatología (Mexico City, Mexico). We also thank all the mothers and babies who donated samples for this study. Work in the SS laboratory was supported by recurrent funding from the Institut National de la Santé et de la Recherche Médicale (INSERM), Aix-Marseille University, and the Ligue Contre le Cancer (Equipe Labellisée 2023). Work in the ORJ lab was supported by the SECIHTI grant CF2019 1727995. CJVM and ACB received PhD grants (960641 and 1136695, respectively) from SECIHTI. LAKC receives a Postdoc grant from SECIHTI (BP-PM-20250513122652290-11414813).

## Data availability statement

The anonymized RNA-seq datasets are deposited in the GEO repository GSE322854, and will be available upon publication.

## Authorship contribution statement

CJVM and LAKC participated in the investigation and formal analysis, wrote the original manuscript, and revised the final version. SMG, ACB, IYCC, and SVR participated in the investigation. ACHR, CI, and SS revised the manuscript. AS participated in the conceptualization of the idea, oversaw the project, wrote the original manuscript, edited, and revised the final version. ORJ conceptualized the idea, oversaw the project, wrote the original manuscript, edited and revised the final version.

---

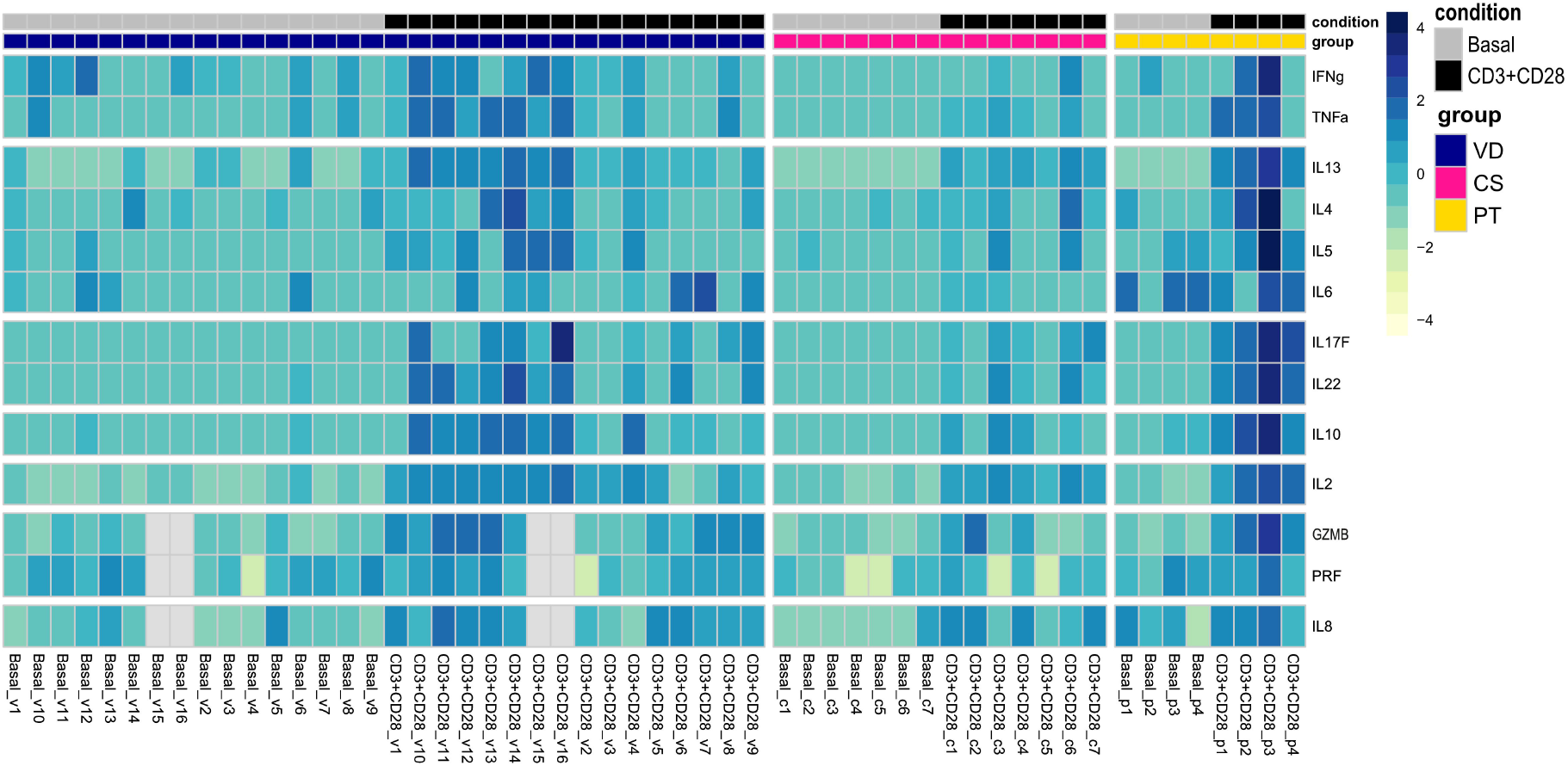

