## Supplementary material for "PREMATURE BIRTH AND CESAREAN SECTION ARE ASSOCIATED WITH SIGNIFICANT DIFFERENCES IN NEONATAL CD4^+^ T CELL GENE EXPRESSION AND FUNCTION": Supp Figures and tables

### Supplementary tables

**Supplementary Table 1. Clinical characteristics of the neonatal cohort.** Clinical characteristics of the neonates included in this study. Mononuclear cells (CBMCs) or CD4+ T cells were obtained from cord blood to perform transcriptomic analysis or flow cytometry experiments. The table summarizes gestational age, sex, birth weight, and the study group assigned to each participant.

**Supplementary Table 2. Differentially expressed genes between cesarean section and vaginal delivery neonates.** List of differentially expressed genes (DEGs) identified in the comparison between CD4+ T cells from cesarean section (CS) and vaginal delivery (VD) neonates, which was obtained using DESeq2. Only genes with an absolute log2 fold change ≥ 1 and adjusted p-value < 0.05 are included. The table reports Ensembl gene identifiers, normalized mean expression (baseMean), log2 fold change, standard error (lfcSE), Wald statistic, P-value, Benjamini–Hochberg adjusted P-value (padj), Entrez Gene ID, HGNC gene symbol, and gene annotation.

**Supplementary Table 3. Differentially expressed genes between preterm and cesarean section neonates.** List of differentially expressed genes (DEGs) identified in the comparison between CD4+ T cells from preterm (PT) and full-term cesarean section (CS) neonates, which was obtained using DESeq2. Only genes with an absolute log2 fold change ≥ 1 and adjusted p-value < 0.05 are presented. Reported variables include Ensembl gene ID, normalized mean expression (baseMean), log2 fold change, standard error (lfcSE), Wald statistic, P-value, Benjamini–Hochberg adjusted P-value (padj), Entrez Gene ID, HGNC gene symbol, and functional gene description.

**Supplementary Table 4. Complete list of differentially expressed genes between cesarean section and vaginal delivery neonates.** Complete results of the differential expression analysis comparing cesarean section (CS) and vaginal delivery (VD) neonates generated using DESeq2. The table includes all analyzed genes together with normalized mean expression (baseMean), log2 fold change, standard error (lfcSE), Wald statistic, P-value, Benjamini–Hochberg adjusted P-value (padj), Entrez Gene ID, HGNC gene symbol, and gene description, irrespective of fold-change threshold.

**Supplementary Table 5. Complete list of differentially expressed genes between preterm and cesarean section neonates.** Complete differential expression analysis comparing preterm (PT) and cesarean section (CS) neonates obtained using DESeq2. The table includes all analyzed genes together with normalized mean expression (baseMean), log2 fold change, standard error (lfcSE), Wald statistic, P-value, Benjamini–Hochberg adjusted P-value (padj), Entrez Gene ID, HGNC gene symbol, and functional annotation, regardless of the magnitude of differential expression.

### Supplementary figures

#### 1) Representative figures of the purity of isolated CD4+ T cells

**A)** Sample purity

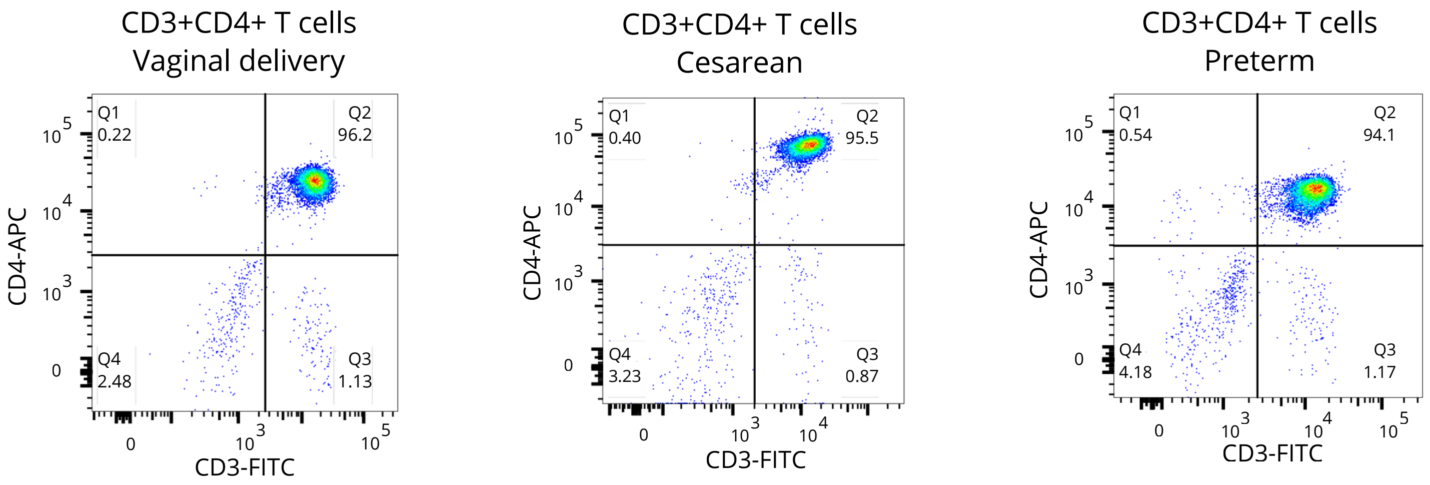


**B)** Percentage of purity

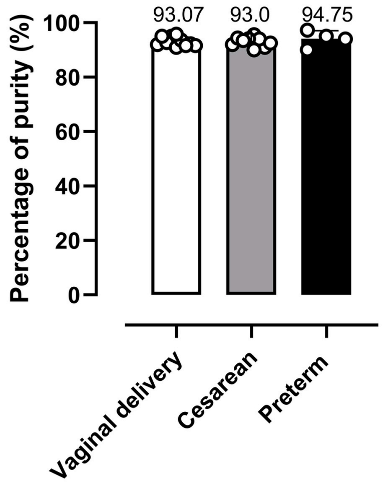


**C)** CD45RO and CD69 expression

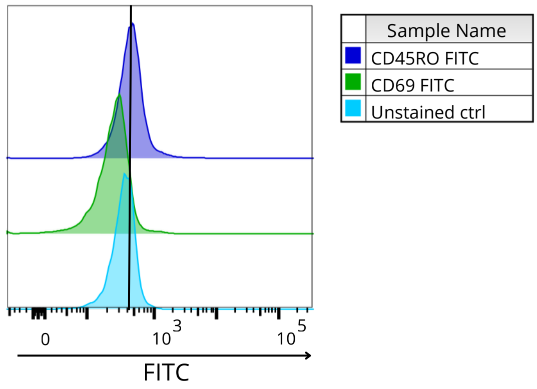


**Supplementary Figure 1:** Purity of isolated CD4+ T cells. **(A)** Representative purity of neonatal samples with different mode of birth and gestational age. **(B)** Mean of percentage of purity (vaginal delivery =16, cesarean = 9, preterm = 4). **(C)** Representative histogram of CD45RO and CD69 expression on neonatal CD4+ T cells.

#### 2) Complementary transcriptomic analyses

**A)** Treeplot of KEGG enriched pathways from overexpressed genes in neonatal CD4+ T cells born by vaginal delivery.

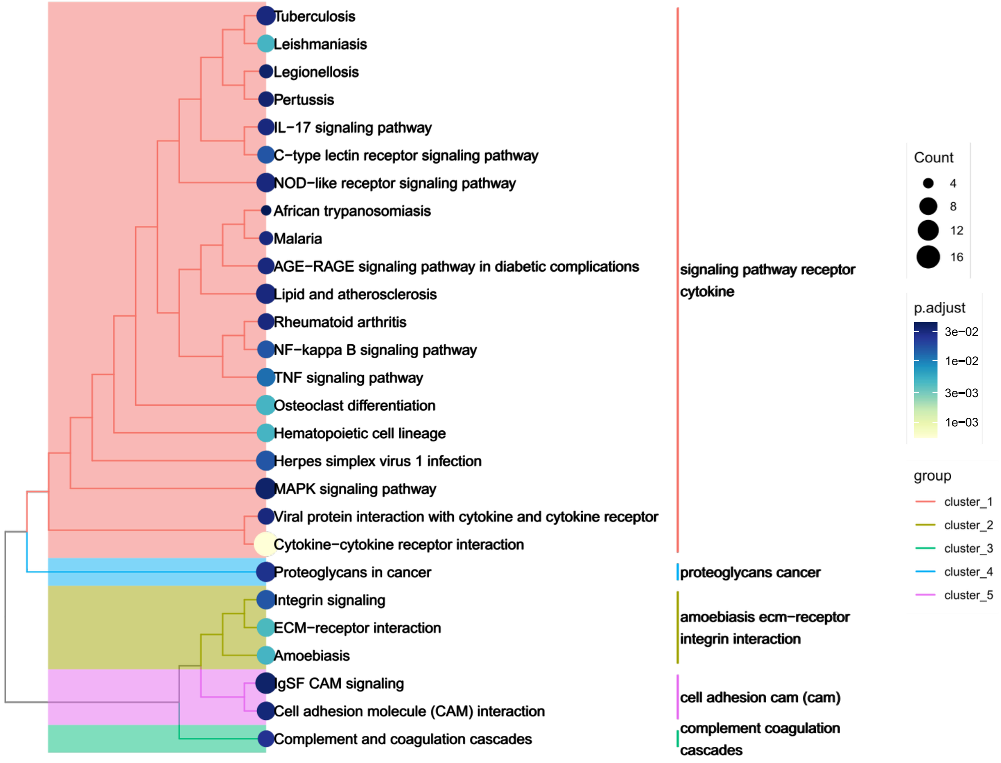


**B)** Volcan plot from differentially expressed genes in neonatal CD4+ T cells born at term or preterm by cesarean section.

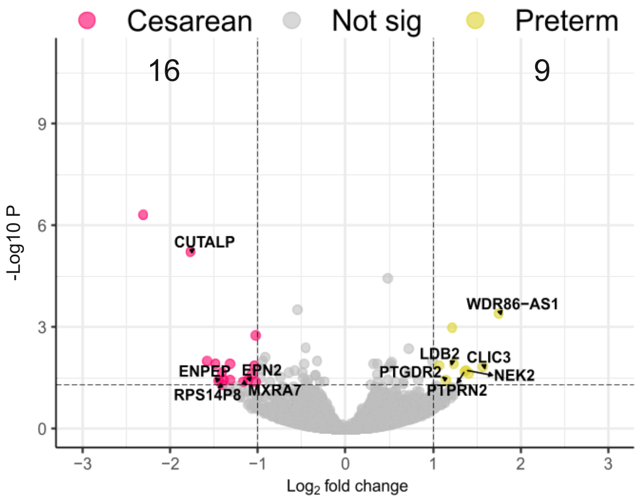


**Supplementary Figure 2:** Complementary transcriptomic analyses. **(A)** Treeplot of KEGG enriched pathways from over expressed genes in neonatal CD4+ T cells born by vaginal delivery. **(B)** Volcanoplot showing the differentially expressed genes between neonatal CD4+ T cells from preterm and cesarean section samples.

### 3) Heatmaps for the differentially expressed genes

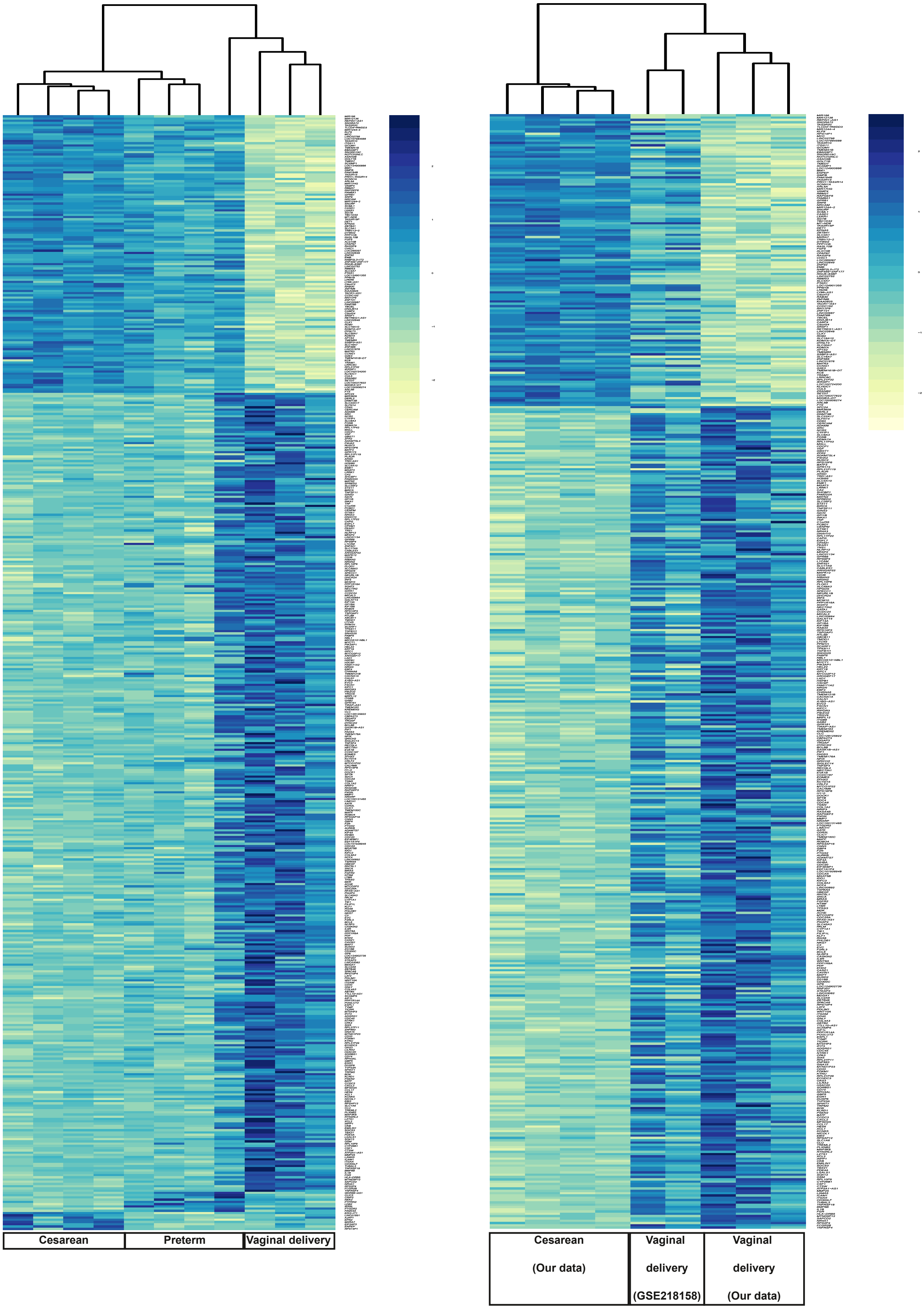


**Supplementary Figure 3.** Heatmap of differentially expressed genes in neonatal CD4+ cells. (A) Heatmap of differentially expressed genes in neonatal CD4+ cells from vaginal delivery, term cesarean section, and preterm cesarean section groups in our cohort. Expression values were normalized and scaled on a per-gene basis (Z-score). (B) Heatmap of the same gene set including two additional samples from a previously published study, generated using the same processing and visualization approach and maintaining gene order to enable direct comparison of the cesarean section versus vaginal delivery group.

#### 4) Complementary gene set enrichment analysis

**A)** Enriched KEGG pathways in CD4+ T cells from term neonates delivered by vaginal delivery or cesarean section.

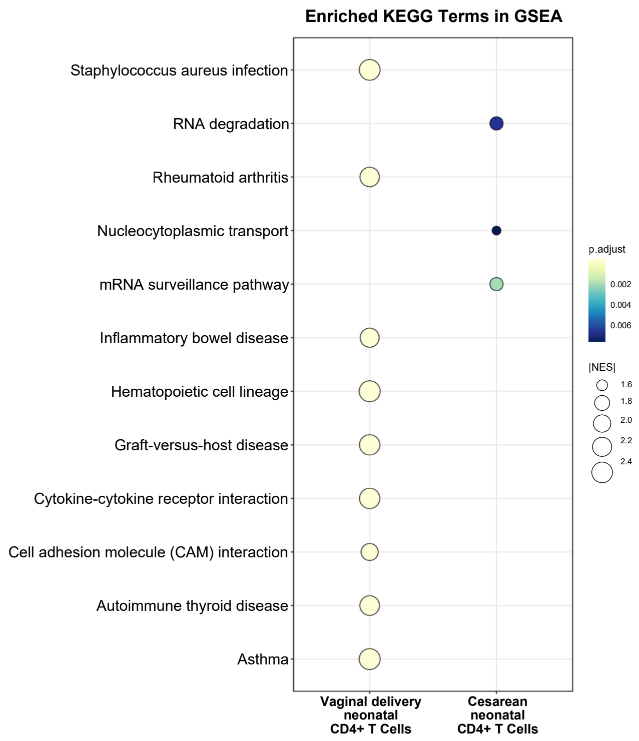


**B)** Enriched KEGG pathways in CD4+ T cells from preterm neonates.

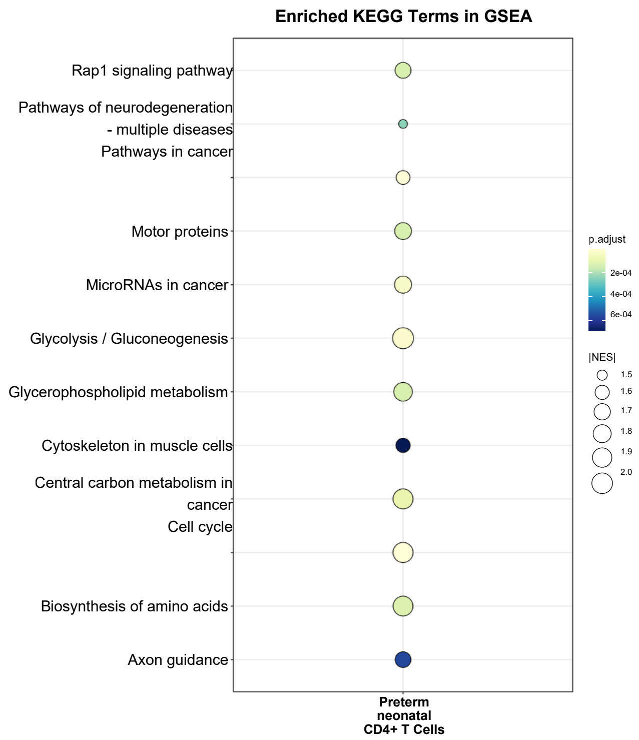


**Supplementary Figure 4:** Enriched KEGG pathways in **(A)** term neonates delivered vaginally or by cesarean section, and **(B)** preterm neonates.

### 5) Gating strategy

#### **A)** Transcription factors gating

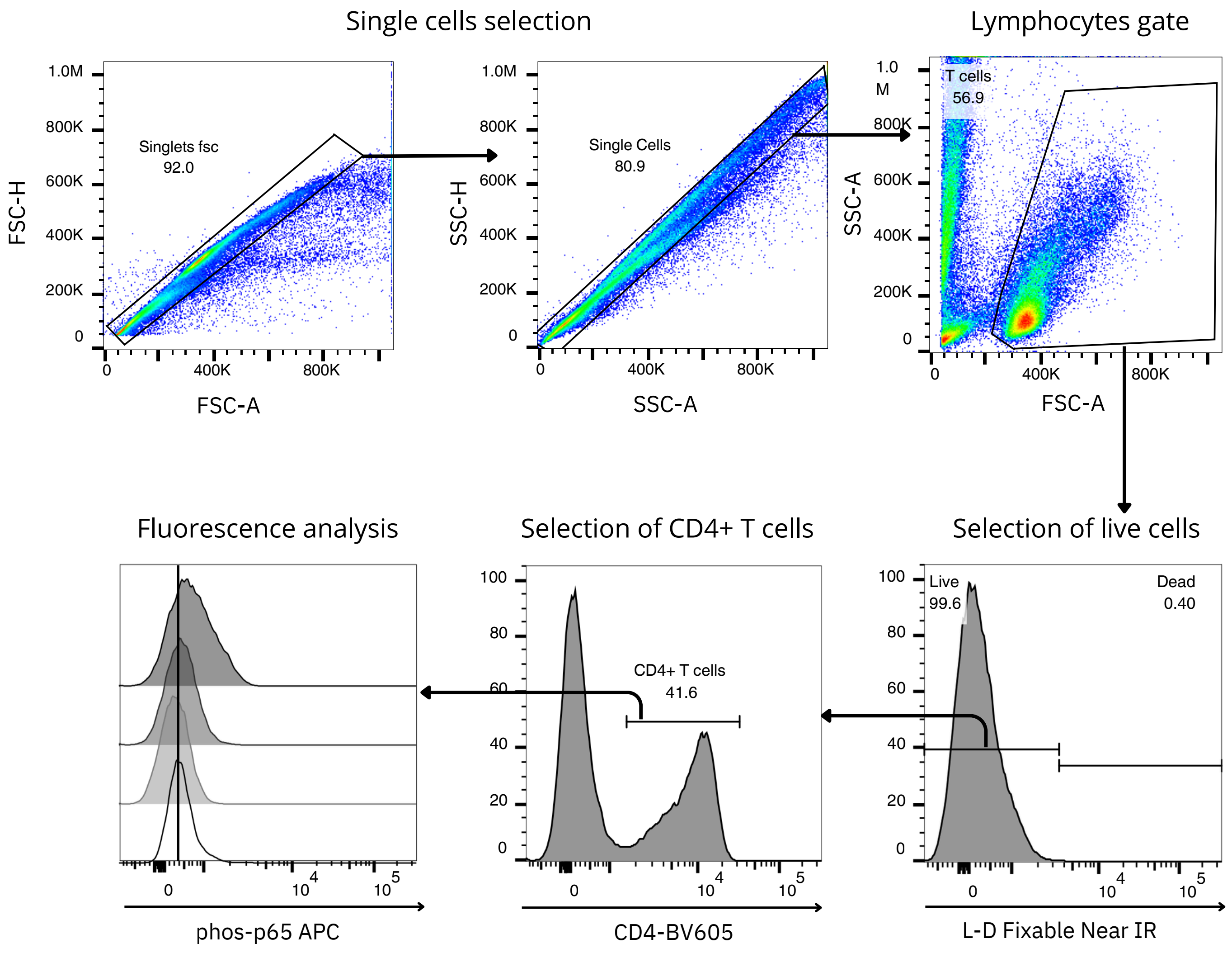

#### B) Proliferation gating

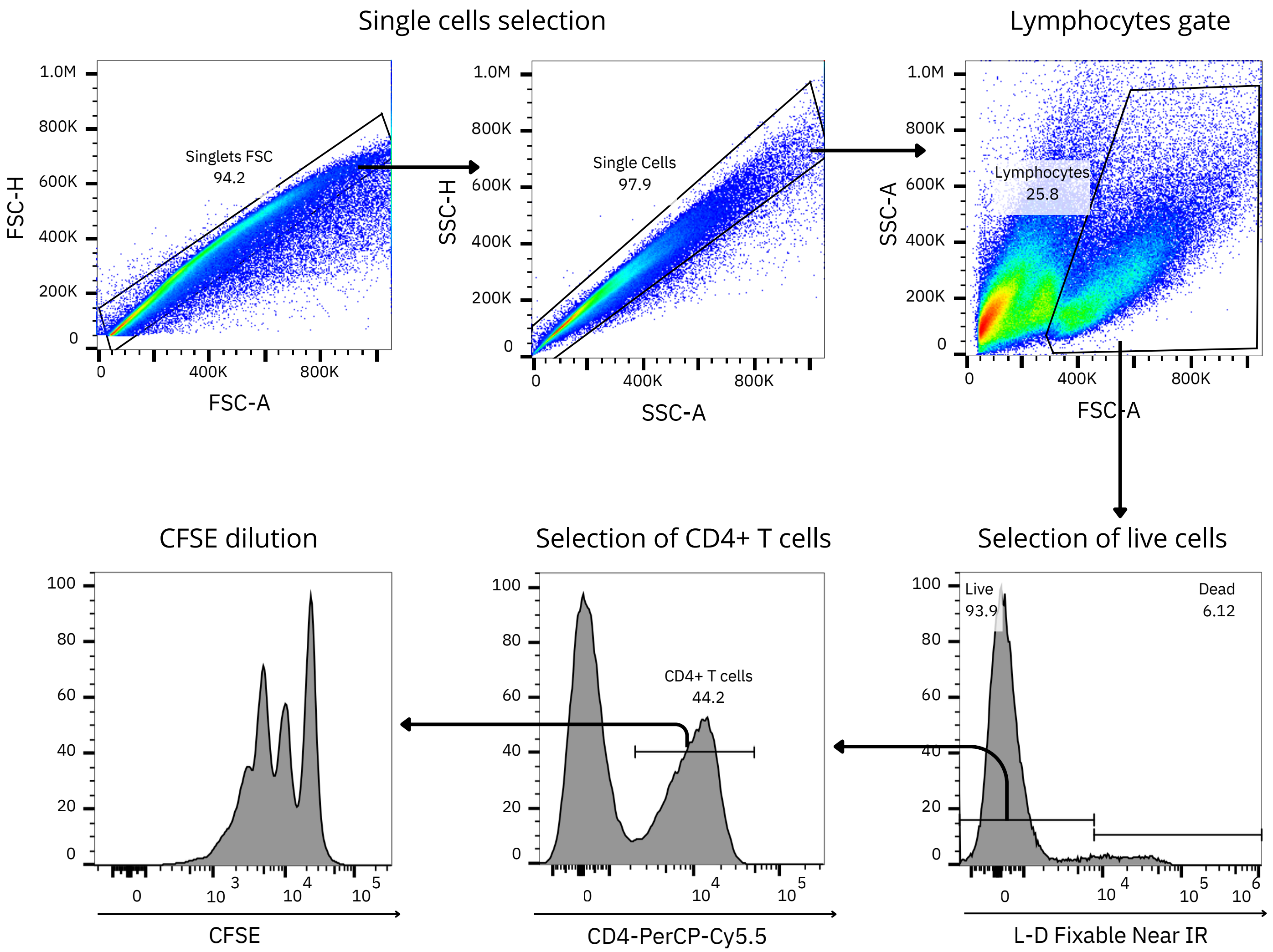

#### C) Fluorescence minus one (FMO) controls

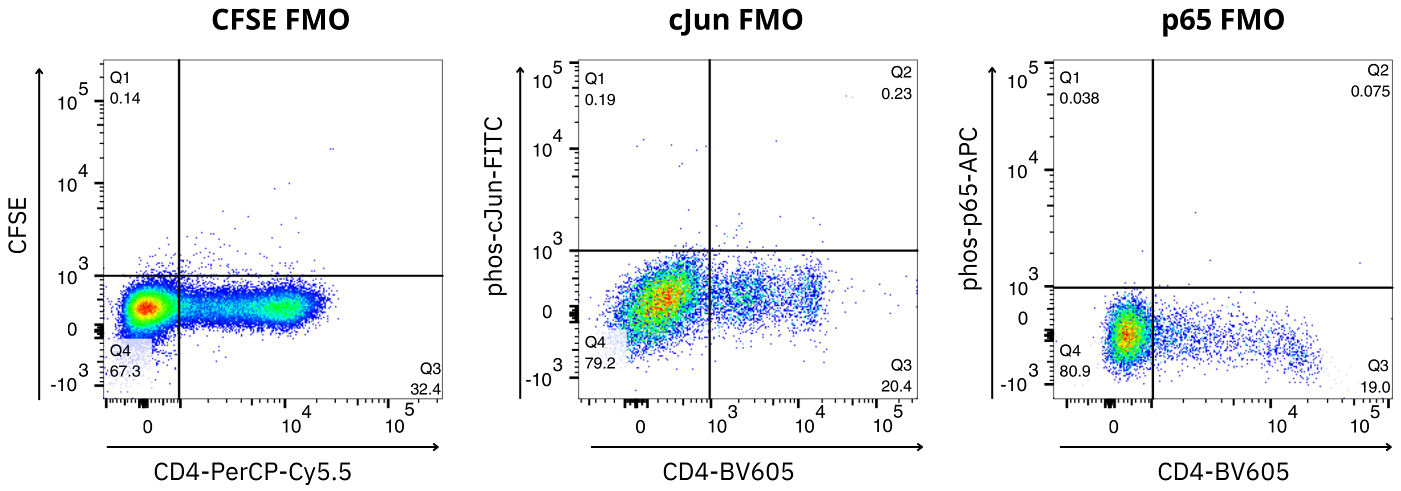


| | Staining |
|---|---|
| Live-dead Near IR | + |
| CD4-PerCP-Cy5.5 | + |
| CFSE | - |

| | Staining |
|---|---|
| Live-dead Near IR | + |
| CD4-BV605 | + |
| phos-cJun-FITC | - |

| | Staining |
|---|---|
| Live-dead Near IR | + |
| CD4-BV605 | + |
| phos-p65-APC | - |

**Supplementary Figure 5:** Gating strategy. **(A)** Gating strategy for transcription factors analysis. **(B)** Gating strategy for proliferation analysis. **(C)** FMO controls for proliferation and transcription factors analysis.

#### 6) Cytokine production by Cord Blood Mononuclear Cells (CBMC)

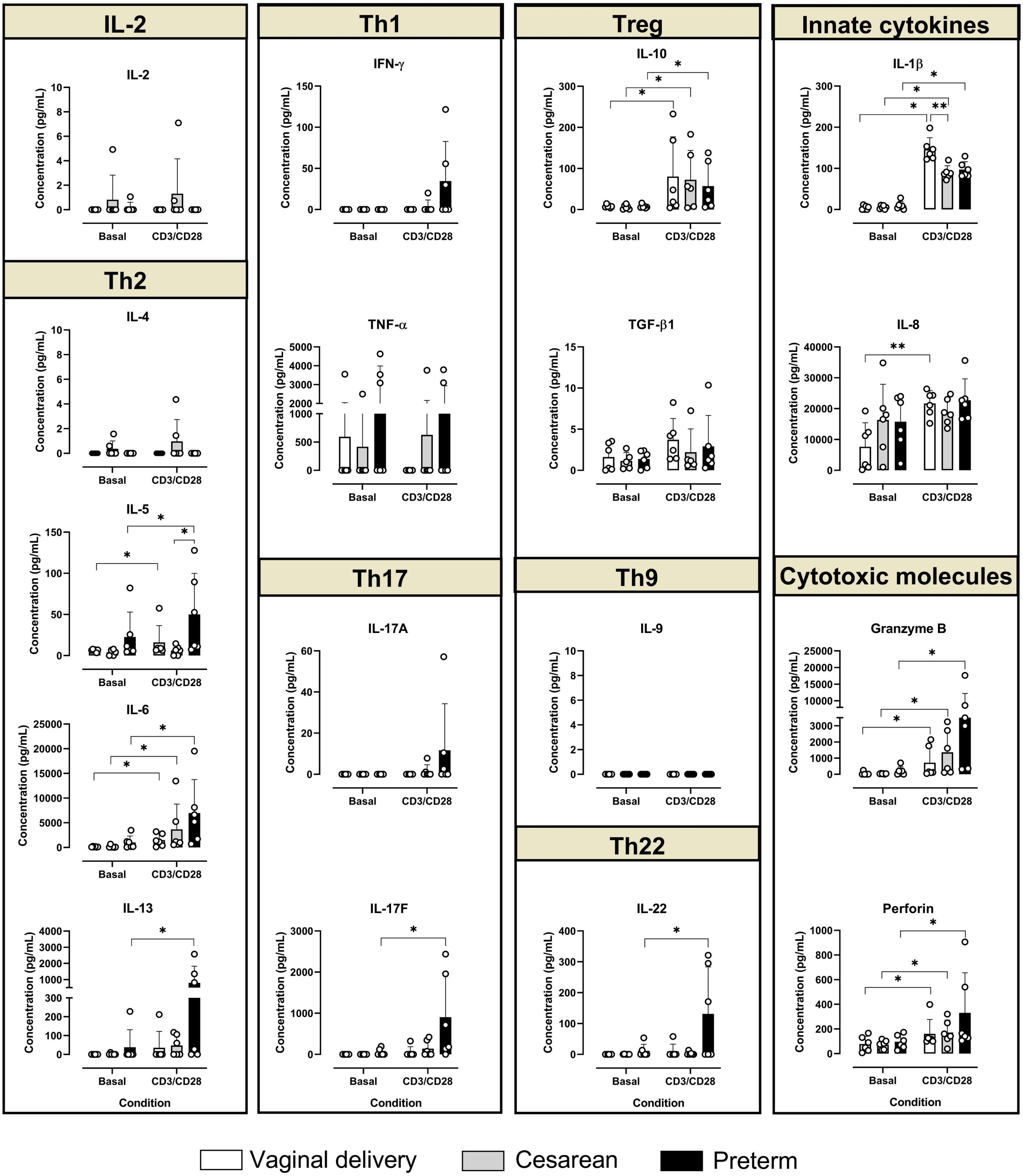


**Supplementary Figure 6.** Cytokine production by cord blood mononuclear cells (CBMC). T helper cytokines, innate cytokines, and cytotoxic molecules produced by CBMC under basal or stimulated conditions after 96 h of incubation. Cytokine concentrations were analyzed using the Wilcoxon test for paired samples and the Kruskal–Wallis test for unpaired samples, followed by the original Benjamini–Hochberg (BH) false discovery rate (FDR) correction for multiple comparisons. $P < 0.05$, $P < 0.01$, $*P < 0.001$. The number of samples was: vaginal delivery ($n = 6$), cesarean section ($n = 6$), and preterm ($n = 6$).
